# SARM1 activation induces reversible mitochondrial dysfunction and can be prevented in human neurons by antisense oligonucleotides

**DOI:** 10.1101/2024.04.02.587827

**Authors:** Andrea Loreto, Kaitlyn M. L. Cramb, Lucy A. McDermott, Christina Antoniou, Ilenia Cirilli, Maria Claudia Caiazza, Elisa Merlini, Peter Arthur-Farraj, Elliot D. Mock, Hien T. Zhao, David L. Bennett, Giuseppe Orsomando, Michael P. Coleman, Richard Wade-Martins

**Author notes:** Lead contact: Dr Andrea Loreto. Correspondence to: Dr Andrea Loreto, Prof Michael Coleman, Prof Richard Wade-Martins. These authors contributed equally to this work (Andrea Loreto, Kaitlyn M. L. Cramb).

## Abstract

SARM1 is a key regulator of a conserved program of axon degeneration increasingly linked to human neurodegenerative diseases. Pathological SARM1 activation causes rapid NAD consumption, disrupting cellular homeostasis and leading to axon degeneration. In this study, we develop antisense oligonucleotides targeting human SARM1, demonstrating robust neuroprotection against morphological, metabolic, and mitochondrial impairment in human iPSC-derived dopamine neurons induced by the lethal neurotoxin vacor, a potent SARM1 activator. Furthermore, our findings reveal that axon fragmentation can be prevented, and mitochondrial dysfunction reversed using the NAD precursor nicotinamide, a form of vitamin B_3_, even after SARM1 activation has occurred, when neurons are already unhealthy. This research identifies ASOs as a promising therapeutic strategy to block SARM1, and provides an extensive characterisation and further mechanistic insights that demonstrate the reversibility of SARM1 toxicity in human neurons. It also identifies the SARM1 activator vacor as a specific and reversible neuroablative agent in human neurons.

## Introduction

Sterile alpha and TIR motif-containing protein 1 (SARM1) is a central executor of programmed axon death, an evolutionarily conserved pathway of axon degeneration that is increasingly associated with human neurodegenerative diseases^1,2^. SARM1 activity in neurons is tightly regulated by the axonal survival enzyme nicotinamide mononucleotide adenylyltransferase 2 (NMNAT2)^3,4^. Both SARM1 and NMNAT2 play a pivotal role in nicotinamide adenine dinucleotide (NAD) metabolism. Depletion or inactivation of NMNAT2 leads to an accumulation of its substrate nicotinamide mononucleotide (NMN), which then binds to SARM1, activating it^5–10^. Once activated, SARM1 rapidly consumes NAD and causes axon degeneration^11,12^. Vacor, a deadly environmental neurotoxin for humans^13^, functions in a similar manner. Vacor undergoes conversion within cells to vacor mononucleotide (VMN)^14^, an analog of NMN, which binds to and potently activates SARM1, ultimately causing neuron death^15^ (Fig. 1A). Other neurotoxins are known to exhibit similar behaviour^16,17^.

**Fig. 1.**
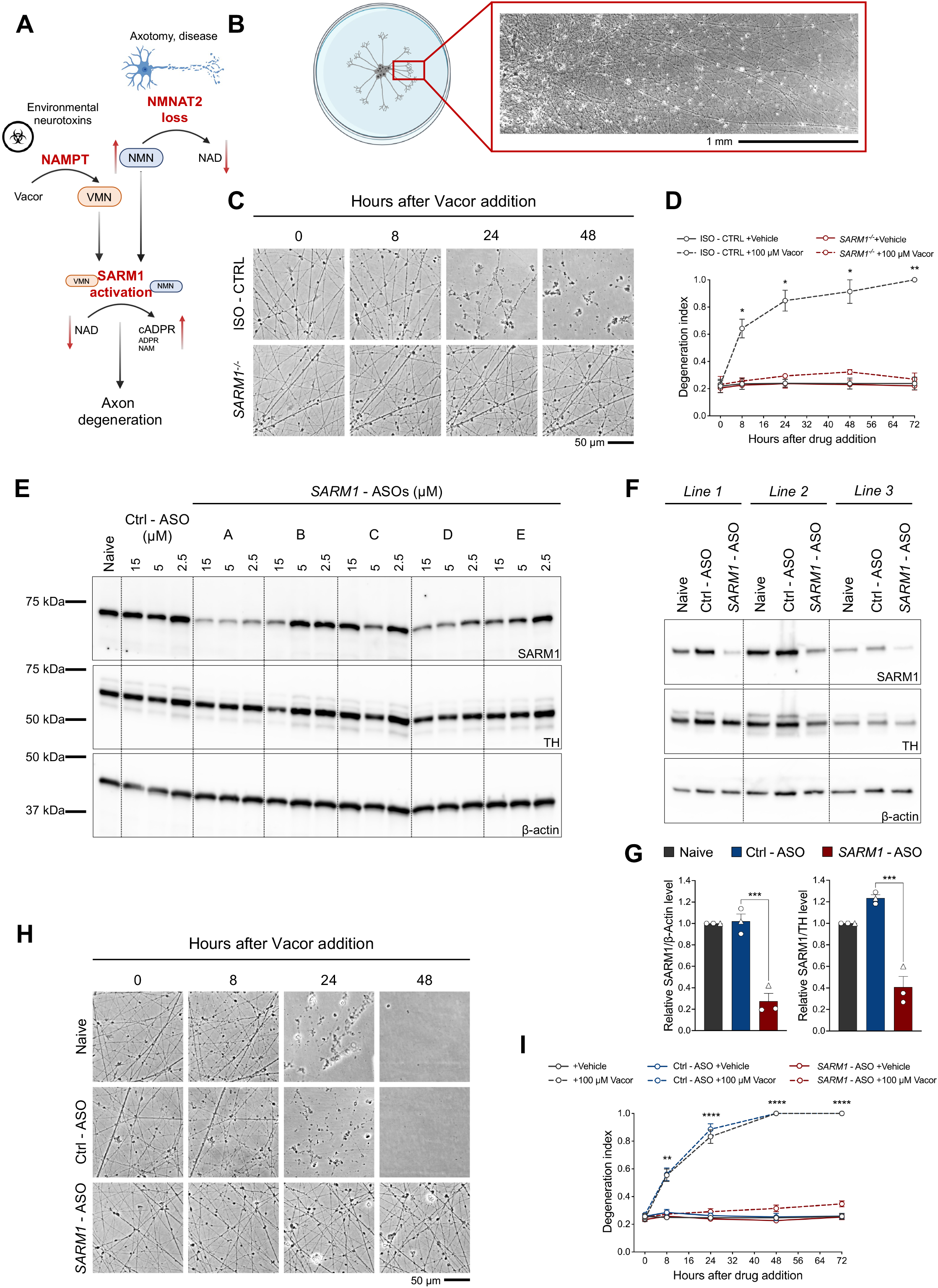
ASOs targeting human *SARM1* rescue human iPSC-derived dopamine neurons from vacor toxicity. **(A)** Schematic overview of programmed axon death and the mechanism by which vacor acts on it. **(B)** Representative image of hiPSC-DANs plated as spot cultures growing long axons. **(C)** Representative images of axons from *SARM1^-/-^* and isogenic control (ISO - CTRL) hiPSC-DANs treated with vacor or vehicle. **(D)** Quantification of the degeneration index for the conditions described in part (C) (mean ± SEM; n = 3 from 3 independent differentiations; two-way RM ANOVA followed by Tukey’s multiple comparison test; statistical significance shown relative to *SARM1^-/-^*+100 µM Vacor). **(E)** Representative immunoblots of hiPSC-DANs untreated (naive) or treated with *SARM1* - ASOs or a non-targeting control ASO (Ctrl - ASO) at the indicated concentrations, probed for SARM1, TH and β-actin (loading controls). **(F)** Representative immunoblots of hiPSC-DANs from 3 healthy individuals untreated, or treated with *SARM1* ‘A’ - ASO (this ASO was employed in all subsequent experiments at a concentration of 5 µM) or a non-targeting control ASO, probed for SARM1, TH and β-actin (loading controls) (*Line 1 = 067; Line 2 = 156; Line 3 = 053*). **(G)** Quantification of normalised SARM1 level (to β- actin and TH) is shown for the conditions described in part (F) (mean ± SEM; n = 3 from 1 independent differentiation; ordinary one-way ANOVA followed by Tukey’s multiple comparison test). **(H)** Representative images of axons from hiPSC-DANs untreated or treated with *SARM1* - ASO and Ctrl - ASO following administration of vacor or vehicle. **(I)** Quantification of the degeneration index for the conditions described in part (H) (mean ± SEM; n = 9 from 3 independent differentiations; two-way RM ANOVA followed by Tukey’s multiple comparison test; statistical comparison shown is Ctrl - ASO +100 µM Vacor vs *SARM1* - ASO +100 µM Vacor).

Targeting programmed axon death holds significant potential for clinical applications^18^. The deletion of SARM1 is remarkably neuroprotective, as evidenced by the complete morphological and functional rescue of neurons in both genetic^19^ and toxic^15^ mouse models of neurodegeneration, each with human equivalents^13,20^. Several studies in animal models suggest the involvement of the programmed axon death pathway across a range of neurodegenerative disorders^2^. Notably, recent research has expanded upon findings derived from animal studies, identifying harmful gene variants of SARM1 and NMNAT2 in human disease. For instance, loss-of-function mutations in NMNAT2 have been observed in patients with varying degrees of severity of polyneuropathy^20–22^. Genome-wide association studies (GWAS) have associated SARM1 with Amyotrophic Lateral Sclerosis (ALS)^23,24^, and gain-of-function mutations in SARM1 are notably prevalent in individuals affected by ALS and other motor nerve disorders^25,26^.

To date, drug development has focused on inhibiting SARM1 activity through the development of small molecule inhibitors and gene therapy approaches^27–30^. However, current preclinical data suggest that SARM1 activity and/or levels must be substantially reduced to achieve strong neuroprotection^31^ and, once activated, SARM1 can rapidly cause axon degeneration. It is therefore important to develop a range of therapeutic strategies targeting SARM1, which can be employed either individually or in combination, to achieve the most effective therapeutic window and optimal neuroprotection. Among these approaches, antisense oligonucleotide (ASO) therapies hold significant promise. ASO-based treatments have already shown remarkable therapeutic effects in clinical settings for neurodegenerative disorders^32^. Their mechanism of action is different from the currently in-development approaches targeting SARM1, as they lead to long-term reduction of the levels of the protein of interest rather than blocking its enzymatic activity^33^. We and others have recently demonstrated that ASOs against mouse SARM1 in rodent neurons confer neuroprotection^31,34^. However, data in human neurons with ASOs targeting human SARM1 are currently lacking.

In addition to developing effective therapies, essential questions remain about the mechanism of programmed axon death. A deeper understanding of the molecular processes involved has revealed specific biomarkers of SARM1 activation and metabolic changes that occur before, or even in the absence, of axon degeneration and neuron loss^8,35–37^. Notably, SARM1 activation alters NAD metabolism in neurons, including a lowering of NAD levels and cADPR accumulation, and how these metabolic changes affect cellular functionality is unclear. Of particular interest is the connection between mitochondria and programmed axon death. We and others have shown that mitochondrial dysfunction can both trigger and result from programmed axon death activation^7,38–40^. Nevertheless, It remains unclear whether potent SARM1 activation marks a critical point from which axons are inevitably committed to degeneration, with the subsequent mitochondrial dysfunction following SARM1 activation serving merely as a marker of degenerating neurons^40^. Linked to the concept of commitment to degeneration is the question of the reversibility of SARM1 toxicity and the associated functional alterations. For example, it is unclear whether morphological damage to axons and mitochondrial dysfunction downstream of SARM1 can be halted or even reversed once SARM1 activation has taken place. Understanding this is important not only for advancing our comprehension of programmed axon death but also for potential clinical applications of therapies attempting to inhibit SARM1, considering that most diseases are diagnosed after pathological processes have been underway for some time. The question of reversibility is also relevant to therapies aiming to activate SARM1 as a selective neuroablative agent to target conditions such as spasticity, dystonia and neuropathic pain, because permanent SARM1 activation in these conditions could lead to unwanted effects and lasting disability.

Lastly, while programmed axon death has been extensively studied in animal models and rodent primary neuronal cultures, there is shortage of data from human neurons. Bridging this gap is essential as we progress toward clinical translation.

Here, we have developed ASOs targeting human SARM1 and assessed their neuroprotective effects in human iPSC-derived dopamine neurons (hiPSC-DANs). Our findings reveal remarkable neuroprotection against morphological damage, metabolic changes, and mitochondrial dysfunction induced by the lethal neurotoxin vacor, which is the most potent SARM1 activator known to date. Furthermore, we demonstrate that morphological damage to axons can be prevented, and mitochondrial dysfunction reversed using nicotinamide (NAM), a form of vitamin B_3_ and an NAD precursor, after SARM1 activation has already occurred. This study indicates that toxicity caused by SARM1 activation in human neurons is reversible. The substantial neuroprotection achieved with *SARM1*-targeted ASOs in human neurons positions them as a promising therapeutic strategy to block or reverse morphological and functional damage caused by programmed axon death activation. These findings also reveal the viability of small molecule SARM1 activators, such as vacor and its metabolite VMN, as selective and reversible neuroablative treatments in human neurons for neurological conditions such as spasticity, dystonia and neuropathic pain.

## Experimental procedures

### Human iPSC-derived dopamine neuron production

Human induced pluripotent stem cells (iPSCs) used in this study (Table 1) originated from the Oxford Parkinson’s Disease Center (OPDC) Discovery cohort. SARM1 - deficient iPSCs were generated using CRIPSR/Cas9 technology. Human iPSC-derived dopamine neurons were produced according to Protocols.io^41^ as previously published^42^. In brief, human iPSCs were plated on Matrigel (BD Biosciences) and expanded in mTeSR1 medium (StemCell technologies) supplemented with mTeSR1 supplement (StemCell Technologies) and 100 U/ml Penicillin and 100 µg/ml Streptomycin (Life Technologies). Cells were passaged using EDTA or TrypLE (Life Technologies) and split 1:2 with 10 µM ROCK inhibitor (Y-27632) (Tocris Bioscience). Cells were differentiated into dopamine neurons using on a modified Krik’s protocol^43^, incorporating modifications adapted from^44^ as previously described^42^. iPSCs were patterned to become ventral midbrain neural progenitors, expanded at this stage before being differentiated into dopamine neurons. hiPSC-DANs were quality controlled for differentiation efficiencies (Fig. S1D,E), half of the media was changed every 2-3 days and all experiments were performed between DIV 35 and 45. For spot cultures, 12-well plates were coated overnight with 1% GelTrex (Thermo Fisher Scientific) diluted in H_2_O. The following day, GelTrex was removed and the well left to dry completely. This allowed to plate 10 µl ‘spots’ containing 100 000 hiPSC-DANs in culturing media at the centre of the well, minimising cell dispersion. Cells were given 30 minutes to attach in the incubator before the well was filled with 1 ml of culturing media.

**Table 1.**
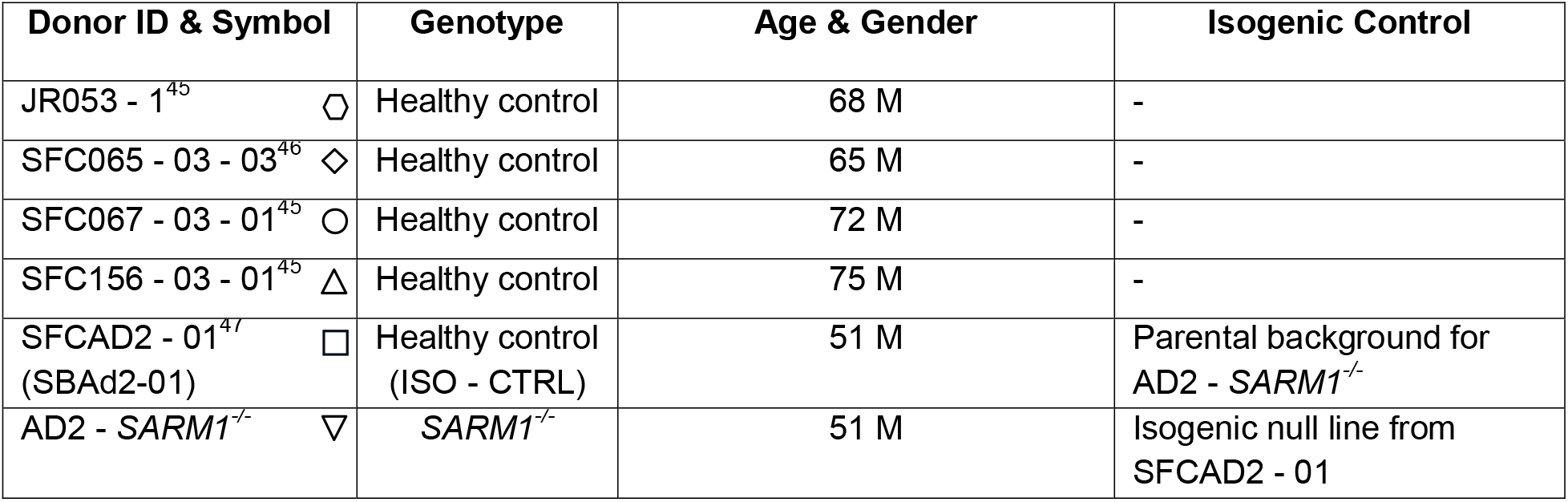
Human iPSC lines used in this study.

### Antisense oligonucleotides

The antisense oligonucleotides (ASOs) were developed and synthesised by Ionis Pharmaceuticals, as previously described^48^. The ASOs are 20 nucleotides in length, chemically modified MOE-gapmer oligonucleotides. The central gap segment comprises ten 2′-deoxyribonucleotides that are flanked on the 5′ and 3′ by five 2′MOE modified nucleotides. Internucleotide linkages are phosphorothioate, and all cytosine residues are 5′-methylcytosines. The sequences of the 5 ASOs targeting human *SARM1* are: ‘A’ - ASO 5′- CTACTTGTTTGTTAGTTCCA -3′; ‘B’ - ASO 5’- TCCACTATTTTTCCCTACCT -3’, ‘C’ - ASO 5′- GTTGCTTTTCCTGTCATTAG -3′, ‘D’ - ASO 5’- GCACATATTTTATTTGCTAC -3’, ‘E’ - ASO 5’- GCAAGATGTTTGCTTACCTG -3’ and a non-targeting Ctrl - ASO 5′- CCTATAGGACTATCCAGGAA -3′. ASO stock solutions were formulated at 15 mM in PBS (without CaCl_2_ and MgCl_2_) (Merck). ASOs were added to the culture media on DIV 22 and subsequently reapplied at every half media change. Therefore, the ASOs were present in the culture media for a minimum of 13 days before starting the experiments. Unless otherwise indicated, *SARM1* ‘A’ - ASO was employed for most experiments in this study at a final concentration of 5 μM.

### Drug treatments and axotomy

hiPSC-DANs neurons were treated with vacor (Greyhound Chromatography) or vehicle (DMSO) just prior to imaging (time 0 hours) or transected using a scalpel between DIV 35 and 45. When used, FK866 (kind gift of Prof Armando Genazzani, University of Piemonte Orientale) was added at the same time as vacor. When used, NAM was added at the same time or 2 and 4 hours after vacor, as detailed in the results section and in (Fig. 4A; Fig. S5B). The drug concentration used is indicated in the figures and figure legends. Vacor was dissolved in DMSO; quantitation of the dissolved stock was performed spectrophotometrically (L340 nm 17.8 mM^−1^cm^−1^). FK866 was dissolved in DMSO and NAM was dissolved in UltraPure DNase/Rnase-free distilled water (Invitrogen).

**Fig. 2.**
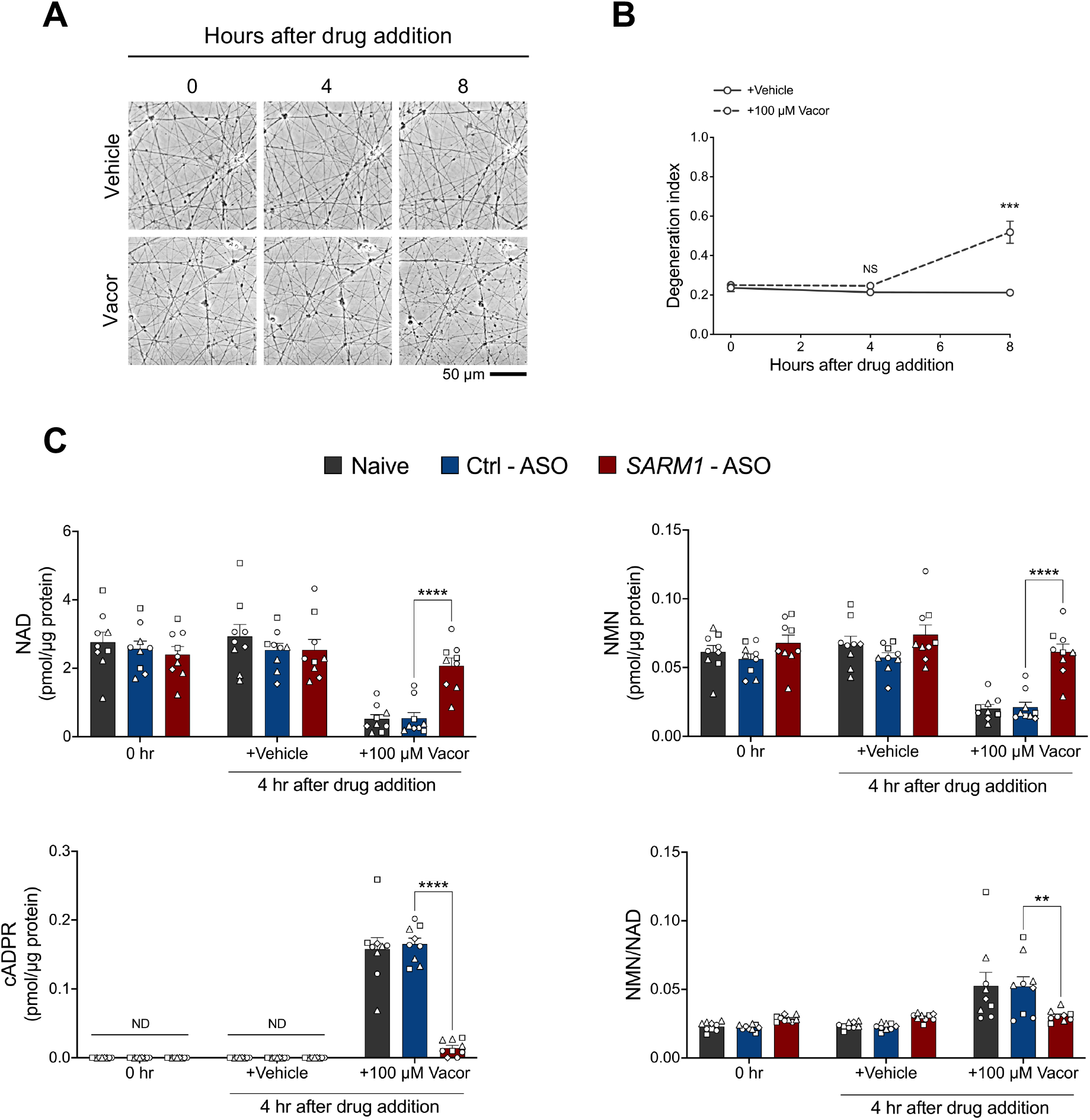
ASOs targeting human *SARM1* prevent metabolic changes caused by SARM1 activation. **(A)** Representative images of axons from hiPSC-DANs at the indicated time points after administration of vacor or vehicle. **(B)** Quantification of the degeneration index for the conditions described in part (A) (mean ± SEM; n = 11 from 3 independent differentiations; two-way RM ANOVA followed by Šídák’s multiple comparisons test). **(C)** NAD, cADPR, NMN levels and NMN/NAD ratio in hiPSC-DANs untreated (0 hr) or treated with *SARM1* - ASO and Ctrl - ASO at the indicated time points following administration of vacor or vehicle (mean ± SEM; n = 9 from 3 independent differentiations; ordinary two-way ANOVA followed by Tukey’s multiple comparison test).

**Fig. 3.**
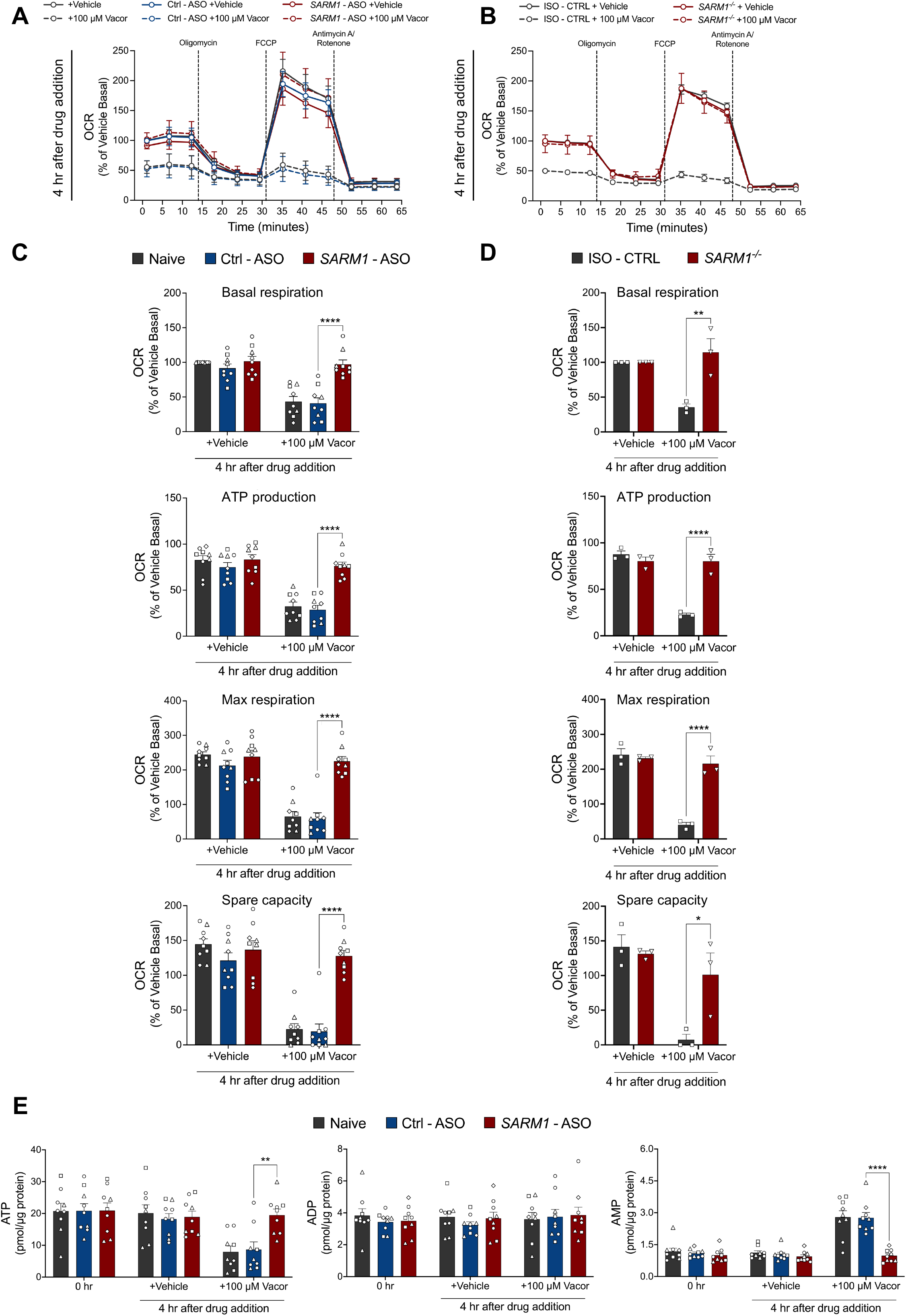
ASOs targeting human *SARM1* prevent mitochondrial dysfunction caused by SARM1 activation. **(A,B)** Mitochondrial respiration in hiPSC-DANs untreated, or treated with *SARM1* - ASO and Ctrl - ASO (A), and in *SARM1^-/-^* and ISO - CTRL hiPSC-DANs (B) at the indicated time point following administration of vacor or vehicle. Oxygen consumption rate (OCR) was normalised to basal respiration of vehicle treated hiPSC-DANs within each individual line and shown as a % change (mean ± SEM; n = 9 from 3 independent differentiations; ordinary two-way ANOVA followed by Tukey’s multiple comparison test). **(C,D)** Quantification of basal respiration, ATP production, maximal respiration and spare capacity for the conditions described in part (A) and (B). **(E)** ATP, ADP and AMP levels in hiPSC-DANs untreated (0 hr), or treated with *SARM1* - ASO and Ctrl - ASO at the indicated time points following administration of vacor or vehicle (mean ± SEM; n = 9 from 3 independent differentiations; ordinary two-way ANOVA followed by Tukey’s multiple comparison test).

**Fig. 4.**
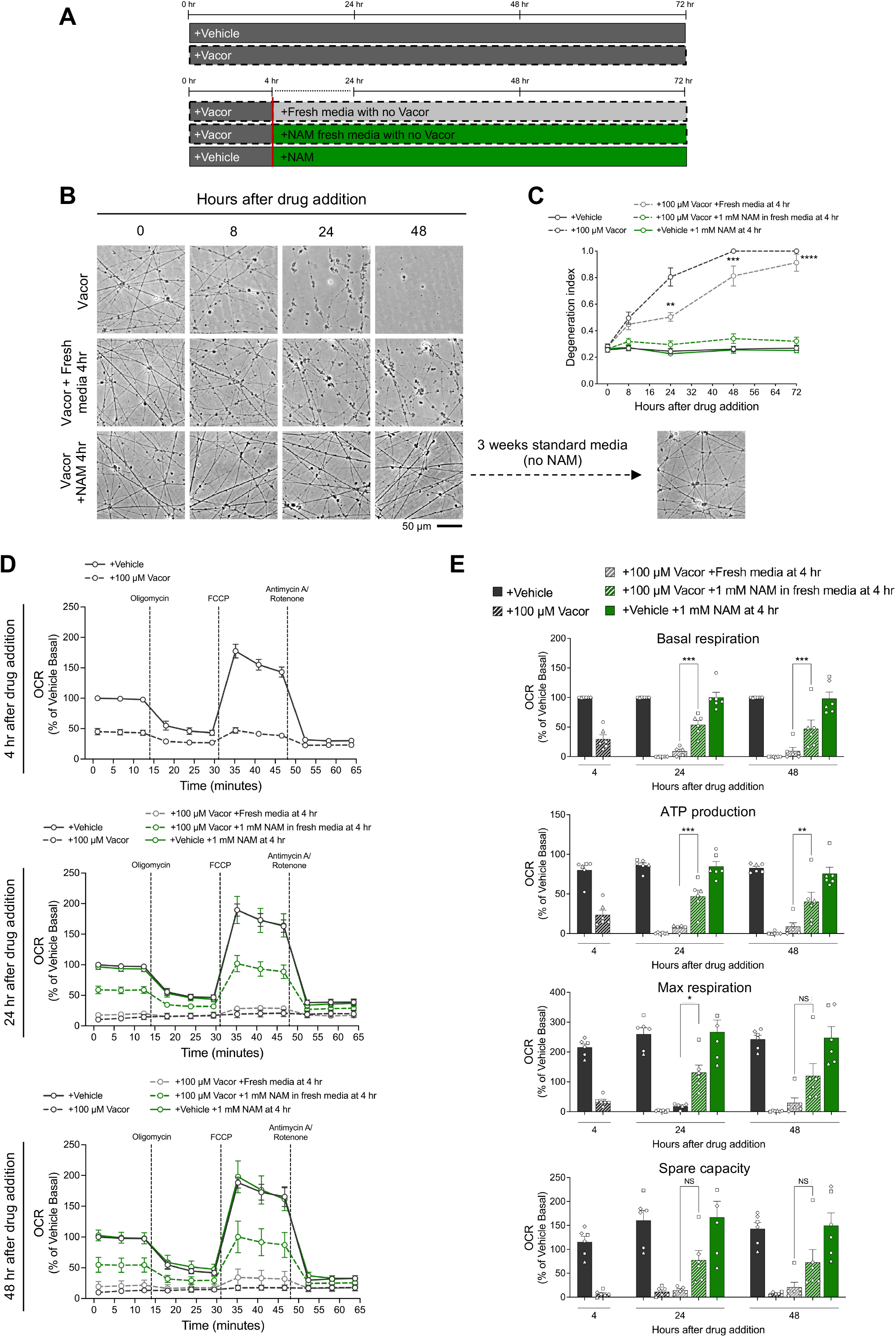
Prevention of axon degeneration and improvement of mitochondrial dysfunction are achievable even after SARM1 activation has occurred. **(A)** Schematic overview of the experimental design for experiments in (B-E). Fresh media with or without NAM was administered 4 hours after vacor treatment while simultaneously removing vacor by replacing culture media. **(B)** Representative images of axons from hiPSC-DANs following treatments outlined in part (A). **(C)** Quantification of the degeneration index for the conditions described in part (B) (mean ± SEM; n = 9 from 3 independent differentiations; Mixed-effects model REML followed by Tukey’s multiple comparison test; statistical comparison shown is +100 µM Vacor +Fresh media at 4 hr vs +100 µM Vacor +1 mM NAM in fresh media at 4 hr). **(D)** Mitochondrial respiration in hiPSC-DANs following treatments outlined in part (A), at the indicated time point following drug administration. OCR was normalised to basal respiration of vehicle treated hiPSC-DANs within each individual line and shown as a % change. **(E)** Quantification of basal respiration, ATP production, maximal respiration and spare capacity for the conditions described in part (D) (mean ± SEM; n = 6 from 2 independent differentiations; ordinary two-way ANOVA followed by Tukey’s multiple comparison test).

### Acquisition of phase contrast images and quantification of axon degeneration

Phase contrast images of hiPSC-DANs axons were acquired on a Nikon ECLIPSE TE2000-E upright fluorescence microscope coupled to a monochrome digital camera. For axon degeneration assays, a 20X objective was used and the degeneration index was determined using a Fiji plugin^49^. For each experiment, the average was calculated from three fields per condition.

### Metabolite analysis

Following treatment with ASOs, vacor and vehicle, hiPSC-DANs were washed in ice-cold PBS and rapidly frozen in dry ice and stored at −80 °C until processed. Tissues were ground in liquid N_2_ and extracted in HClO_4_ by sonication, followed by neutralisation with K_2_CO_3_. AMP, ADP, ATP and cADPR levels were measured by direct UV analysis by ion-pair C18-HPLC in the presence of tetrabutylammonium hydrogen sulfate. NMN and NAD levels were measured by spectrofluorometric HPLC analysis after derivatisation with acetophenone^50^. Metabolite levels were normalised to protein levels quantified with the Bio-Rad Protein Assay (Bio-Rad) on formate-resuspended pellets from the HClO_4_ extraction.

### Seahorse Assay

hiPSC-DANs were plated in a XF96 Polystyrene Cell Culture Microplate (Seahorse Bioscience) at 50,000 cells/ well. Assay medium was prepared fresh the day of the assay using XF Base Medium (Agilent Technologies) with 10 mM Glucose (Merck), 1 mM Sodium Pyruvate (Merck) and 2 mM L-Glutamine (Thermo Fisher Scientific). Mitochondrial respiration was measured using a Seahorse Xfe96 Analyzer (Agilent Technologies). Three baseline recordings were taken followed by three recordings after each of the following subsequent injections: 1 μM Oligomycin (Merck), 1 μM FCCP (Merck) and 5 μM Rotenone and Antimycin (R/A) (Merck). Mitochondrial respiration was measured using a Seahorse Xfe96 Analyzer (Agilent Technologies) and normalised to basal oxygen consumption rate. For each experiment, the average was calculated from three to five wells per condition.

### Western blot

hiPSC-DANs cell pellets were collected in ice-cold PBS and were homogenised in RIPA buffer containing protease (cOmplete EDTA-free, Roche) and phosphatase inhibitors (PhosSTOP, Roche). Protein concentration was determined by BCA assay (Thermofisher). 5 μg of sample were loaded on a 4-to-20% SDS polyacrylamide gel (Bio-Rad). Membranes were blocked for 3 hours in 5% milk in TBS (50 mM Trizma base and 150 mM NaCl, PH 8.3, both Merck) plus 0.05% Tween-20 (TBST) (Merck), incubated overnight with primary antibody in 5% milk in TBST at 4 °C and subsequently washed in TBST and incubated for 1 hours at room temperature with HRP-linked secondary antibody (Bio-Rad) in 5% milk in TBST. Membranes were washed, treated with SuperSignal West Dura Extended Duration Substrate (Thermo Fisher Scientific) and imaged with BioRad imaging system. The following primary antibodies were used: rabbit anti-SARM1 (Cell Signaling Technology, 13022, 1:2000), rabbit anti-tyrosine hydroxylase (TH) (Merck, AB152, 1:30 000) and mouse anti-β-actin (Merck, A5316, 1:50 000) as a loading control. Quantification of band intensity was determined by densitometry using Fiji.

### Immunocytochemistry

hiPSC-DANs were plated on a 96-well plate, were fixed for 7 minutes using 4% PFA, permeabilized with 0.1% Triton-X in PBS, blocked for 1 hour with 10% serum in PBS and incubated overnight at 4LC with the following primary antibodies in PBS: chicken anti-MAP2 (Abcam, ab92434, 1:500), rabbit anti- tyrosine hydroxylase (TH) (Merck, AB152, 1:30 000). Cells were then washed with PBS and incubated for 1 hour at room with the following secondary antibodies: donkey anti-rabbit Alexafluor 488 and donkey anti-chicken Alexafluor 647 (both Thermo Fisher Scientific, A21206, A78952) washed one time with PBS and one time with PBS+DAPI (1:10 000) and subsequently imaged on the Opera Phenix High Content Screening System (Perkin Elmer) on a 40X water objective. Differentiation efficiency was quantified using ImageJ macro ‘cell counter’ for each differentiation using three pictures per well, three wells per line.

### Statistical analysis

Graphs were produced using Prism GraphPad software as was the analysis to test for statistical significance (GraphPad Software, La Jolla, USA). A total of five hiPSC control lines derived from different healthy individuals were used throughout this study. For all experiments, except those involving the *SARM1^-/-^* line and its isogenic control, a minimum of three lines were differentiated in parallel and data originate from multiple, independent differentiations. The hiPSC lines used in each experiment are identifiable by a unique symbol (Table 1). The appropriate statistical test used, the n number (reflecting the total number of hiPSC lines used across multiple differentiations) and the number of differentiations for each experiment are indicated in the figure legends. A p-value < 0.05 was considered significant. In figures, p-value > 0.05 is reported as NS (not significant), p-value between 0.01 and 0.05 is reported as *, between 0.001 and 0.01 is reported as **, between 0.0001 and 0.001 Is reported as ***, and p-value < 0.0001 is reported as ****.

## RESULTS

### ASOs targeting human *SARM1* rescue human iPSC-derived dopamine neurons from vacor toxicity

To assess axon degeneration in human neurons, we employed a spot culture technique to confine the cell bodies of hiPSC-DANs within a small area at the center of the cell culture dish. Fifteen days after the last replating, these spot-cultured hiPSC-DANs had developed long axons radiating outward, which was optimal for subsequent imaging and quantification of axon degeneration (Fig. 1B).

We have previously demonstrated that the neurotoxin vacor specifically activates SARM1 in mouse neurons, resulting in axon degeneration and neuronal soma death^15^. We first sought to establish whether vacor induces axon degeneration in human neurons through the same mechanism. To this end, we generated a *SARM1* knockout (*SARM1^-/-^*) hiPSC line using CRISPR/Cas9 gene editing. We validated the line by confirming the absence of detectable SARM1 protein following differentiation into dopamine neurons (Fig. S1A) and axon protection after axotomy^1,51^ (Fig. S1B,C). *SARM1* deletion did not affect the efficiency of differentiation into dopamine neurons, as evidenced by the percentage of neurons expressing dopamine neuronal markers such as tyrosine hydroxylase (TH), which were comparable to those derived from the isogenic control line (ISO - CTRL) as well as other healthy individuals that were used throughout the study (Fig. S1D,E). We then tested whether vacor induces SARM1- dependent axon degeneration in human neurons. We observed dose-dependent axon degeneration in hiPSC-DANs (Fig. S2A,B) and, notably, complete prevention of vacor-induced axon degeneration was observed in *SARM1^-/-^* hiPSC-DANs (Fig. 1C,D). Consistent with findings in mouse neurons, vacor toxicity was also alleviated by inhibiting nicotinamide phosphoribosyltransferase (NAMPT) with FK866, thereby blocking the conversion of vacor into the toxic metabolite and SARM1 activator VMN^14,15^ (Fig. S2C,D). Thus, vacor requires SARM1 to cause axon degeneration in human neurons.

Next, we designed five ASOs that target human *SARM1* and evaluated their effectiveness in reducing SARM1 levels in hiPSC-DANs. All ASOs reduced SARM1 expression levels with variable efficiencies, as compared to both the non-targeting control ASO (Ctrl - ASO) and untreated hiPSC-DANs (Naive). *SARM1* ‘A’ - ASO demonstrated the most efficient knockdown (Fig. 1E) consistently across hiPSC-DAN lines from different healthy individuals (Fig. 1F,G) and robustly preserved axons from degeneration following vacor treatment (Fig. 1H,I). The strength of protection correlated with the efficiency of knockdown, as the less effective *SARM1* ‘D’ - ASO also provided axonal protection, albeit to a lesser extent (Fig. S3A,B). In contrast, the ineffective *SARM1* ‘C’ - ASO failed to protect axons following vacor administration (Fig. S3C,D). This correlation provides evidence that the protection against axon degeneration is linked to the ASOs’ ability to reduce SARM1 levels and does not result from off-target effects. *SARM1* ‘A’ - ASO was employed in all subsequent experiments. These findings indicate that ASOs targeting SARM1 are a viable approach to block programmed axon death in human neurons.

### ASOs targeting human *SARM1* prevent metabolic changes and mitochondrial dysfunction caused by SARM1 activation

We next aimed to investigate whether ASOs targeting SARM1 could not only rescue morphological damage but also block SARM1-dependent metabolic changes that may impact cellular functionality. Active SARM1 rapidly consumes NAD and produces cADPR before observable axon degeneration and therefore, can serve as a marker specific for SARM1 activity in neurons^15,35^. We first tested if there are metabolic changes in vacor-treated hiPSC-DANs before axon degeneration was observable. We collected hiPSC-DANs 4 hours after vacor treatment, at which point axons appeared morphologically unchanged (Fig. 2A,B), and we measured the metabolite levels. We found a marked decrease in NAD levels, along with an increase in cADPR levels and the NMN/NAD ratio. These metabolic changes were rescued by *SARM1* - ASO (Fig. 2C). Interestingly, and in line with findings in mouse neurons, NMN levels were also lowered in a SARM1-dependent manner^15^ (Fig. 2C).

NAD is a key molecule for mitochondrial energy metabolism and mitochondrial dysfunction has been reported after programmed axon death activation in mouse neurons^40^. We therefore evaluated whether specific SARM1 activation impairs mitochondrial respiration in hiPSC-DANs using the Seahorse Assay, and if this occurs prior to, or simultaneously alongside axon degeneration. In a time-course experiment of basal oxygen consumption rate (OCR), we observed that mitochondrial impairment began approximately 2 hours after vacor treatment, indicating that mitochondrial respiration began to decline in a SARM1-dependent manner well before axon degeneration was detectable (Fig. S4A,B). We found that 4 hours after vacor treatment, when no axon degeneration is observed (Fig. 2A,B) there was a dramatic, SARM1- dependent disruption in mitochondrial respiration resulting in impaired basal respiration, maximal respiration, ATP production, and spare capacity (Fig. 3A-D). Remarkably, mitochondrial respiration remained unaltered in *SARM1* - ASO treated hiPSC-DANs, matching the rescue observed with *SARM1* deletion (Fig. 3A-D). Furthermore, HPLC measurements revealed a substantial loss of ATP, and a concomitant increase in AMP level caused by vacor treatment, while ADP levels remained unchanged (Fig. 3E). This disruption was once again rescued by the application of *SARM1* - ASO (Fig. 3E).

Taken together, these data demonstrate that ASOs targeting SARM1 effectively rescue both morphological axonal damage, metabolic changes and cellular functionality in human neurons following SARM1 activation. They also confirm that SARM1 activation is detectable before morphological damage is present and that mitochondrial dysfunction precedes axon degeneration in hiPSC-DANs after SARM1 activation, and this coincides with detectable metabolic changes.

### Prevention of axon degeneration and improvement of mitochondrial dysfunction are achievable even after SARM1 activation has occurred

The increase in cADPR levels, the marked lowering of NAD levels and mitochondrial dysfunction indicate that SARM1 is active in hiPSC-DANs by 4 hours of exposure to vacor when no observable axon degeneration is evident. The question remains whether SARM1 activation signifies an irreversible commitment to degeneration and whether changes in cellular functionality are, at least in part, a marker of degenerating neurons or an active process that can be reversed.

To address this, we designed a series of experiments to determine if axon degeneration can be rescued and mitochondrial dysfunction reversed once SARM1 activation has already occurred. We have previously shown that NAM protects neurons against vacor toxicity by competing with vacor for NAMPT, thereby reducing the accumulation of the toxic vacor metabolite and the SARM1 activator VMN^15^. Additionally, NAM might also enhance NAD production and counteract NAD depletion downstream of SARM1 activation (Fig. S5A). We therefore hypothesised that NAM treatment could reverse vacor toxicity hours after vacor addition, when metabolite changes and mitochondrial dysfunction were already evident. Initially, we treated hiPSC-DANs with NAM either concurrently with vacor, or 2 and 4 hours after vacor exposure (Fig. S5B). NAM treatment significantly delayed axon degeneration in hiPSC-DANs, whether it was added at the same time of vacor, or hours after (Fig. S5C,D). Although delayed, axon degeneration still occurred under these conditions over the following 72 hours (Fig. S5C,D). To test whether vacor toxicity could be fully reversed, we administered NAM 4 hours after vacor treatment while simultaneously removing vacor by replacing culture media with fresh media containing NAM (Fig. 4A). Remarkably, under these conditions axon degeneration was completely halted (Fig. 4B,C). In contrast, vacor removal and its replacement with fresh media without NAM marginally delayed but did not prevent degeneration (Fig. 4B,C). Three days after treatment with vacor and NAM, we replaced the media and reverted to standard media conditions without extra NAM addition, and the axons remained morphologically intact for the subsequent weeks (Fig. 4B), indicating a complete rescue of the morphological damage to axons.

Next, we sought to determine if mitochondrial dysfunction could be reversed after its onset, following SARM1 activation. As previously demonstrated, severe mitochondrial impairment was evident 4 hours following vacor treatment. Using the same experimental paradigm of adding NAM and removing vacor at 4 hours, we observed a significant improvement in mitochondrial functionality at both 24 and 48 hours (Fig. 4D,E). Multiple mitochondrial parameters, including basal respiration, ATP production and maximal respiration were significantly improved compared to hiPSC-DANs that received fresh media without NAM at the 4-hour mark (Fig. 4D,E). Moreover, we found a similar improving trend in spare capacity, although statistical significance was not reached (Fig. 4D,E)

Taken together, our experiments show that axon degeneration can still be rescued, and mitochondrial dysfunction can be improved, hours after SARM1 activation has taken place when metabolic changes and cellular dysfunction are already observable. These data demonstrate that SARM1 activation does not inevitably lead to axon degeneration, and its detrimental effects on cellular functionality can be reversed.

## Discussion

In this study, we show that ASOs targeting human SARM1 are a promising therapeutic approach to block programmed axon death in human neurons. Our findings indicate that ASOs effectively delay morphological damage to axons and improve cellular functionality, including metabolic changes and mitochondrial dysfunction resulting from SARM1 activation. Notably, ASOs achieve this neuroprotective effect against vacor toxicity, a neurotoxin lethal to humans and currently the most potent known chemical activator of SARM1 and of programmed axon death. These ASOs are also a valuable research tool due to their ease of use, being readily added to cell culture media. This makes them particularly advantageous in human iPSC models, enabling the study of treatment effects on the same cell line potentially reducing variability, a common limitation in iPSC models. Furthermore, mechanistic insights suggest that morphological damage can be prevented, and functional damage can be reversed after SARM1 activation has occurred, which has important implications for clinical translation. Lastly, we provide an extensive characterisation of the consequences of SARM1 activation in human neurons, filling an important gap in the field. As we previously suggested^15^, vacor proves to be an excellent tool for drug discovery and can be used in large-scale drug screening programs targeting SARM1 in human neurons.

In recent years, ASO-based therapies have gained momentum, with several receiving FDA approval, thus highlighting their potential as a safe and effective therapeutic strategy. Notably, some of these therapies have yielded remarkable clinical outcomes^32^. ASOs targeting human SARM1 are part of a broader array of approaches under development to target SARM1, which also include small molecule inhibitors and gene therapy^27–29^. Given that SARM1 can trigger rapid axon degeneration once activated, therapeutic strategies must carefully consider patient compliance with the drug regimen. Combining approaches that achieve long-term reduction of protein levels with inhibition of enzymatic activity could potentially yield optimal clinical outcomes. In addition, having different therapeutic options is important for tailoring treatments to individual patients and maximising benefits while minimising potential risks. This is important because, while targeting SARM1 holds promise as a therapy, the physiological roles of SARM1 are still not fully understood, particularly in the context of potential side effects. Largely positive data from animal studies support the reduction in SARM1 as a favourable therapeutic target as SARM1-deficient animals generally maintain good health even into old age^19^. However, these animals are kept in highly controlled, pathogen-free environments, which could mask potential detrimental effects of SARM1 deficiency. For instance, several studies suggest that SARM1 may have important roles in preventing viral damage or spread^52–54^. Other research also suggests a physiological role of SARM1 in regulating synaptic plasticity^55^ and preventing local inflammation^56^. Depending on the specific disease context and pathogenesis, localised ASO treatments may offer advantages over other putative SARM1-targeting therapies that act systemically, as they can help minimize off-target effects. As the first trial targeting SARM1 is underway, there is a pressing need to address these considerations more directly.

Another significant finding from our study is that SARM1 activation induces dramatic cellular dysfunction, yet it does not signify a point of no return where axons are irreversibly committed to degeneration. We expand on previous research that, using biochemical assays, has suggested that SARM1 enzymatic activity is reversible^8^. While our study does not directly investigate the cellular reversibility of SARM1 enzymatic activity, it does provide evidence that axons can be preserved, and mitochondrial dysfunction can be improved after SARM1 activation is detected by metabolic changes and significant cellular dysfunction has already occurred. This reversibility is a critical aspect for clinical translation, considering that when most neurodegenerative diseases are diagnosed, after symptoms have already set in, cellular dysfunction such as mitochondrial impairment and axon degeneration are well underway^57^. The observed improvement in mitochondrial function suggests that, to some extent, mitochondrial dysfunction is linked to metabolic changes triggered by SARM1 activation, rather than being caused by irreversible organelle damage. The dramatic loss of NAD due to SARM1 activation likely significantly contributes to mitochondrial respiration failure. These findings align with recent studies that demonstrate a role for SARM1 in exacerbating mitochondrial dysfunction in a model of Charcot-Marie-Tooth disease type 2A (CMT2A)^58^, which emphasises the importance of further investigating the impact of SARM1 on mitochondrial function.

Distinct from strategies inhibiting SARM1 to protect neurons, this study also identifies direct SARM1 activators, such as vacor and its metabolite VMN, as viable and reversible neuroablative agents in human neurons. We and others have previously shown that SARM1 activators cause specific neurodegeneration in mouse neurons^15,17^, and it had been postulated by Wu and colleagues that these could be used as neuroablative agents in certain conditions^17^. However, until recently, it had not been known whether SARM1 was exclusively expressed in neurons, which would limit the use of SARM1 activators for selective neuroablation. Interestingly, it has recently been shown that oligodendrocytes, the myelinating glia in the central nervous system, contain high levels of SARM1 protein and are very sensitive to SARM1 activators^59,60^. SARM1 has also been detected in murine astrocytes^61^. However, Schwann cells, the glia of the peripheral nervous system, contain no functionally significant SARM1 protein and are completely insensitive to vacor and other SARM1 activators^59^. The combination of these findings with our data showing that SARM1 activation in human neurons is reversible raises the possibility that highly specific SARM1 activators could be peripherally administered for selective and reversible neuroablation in neurological conditions characterised by dystonia, spasticity, and neuropathic pain.

Lastly, while our current study employs vacor, which induces rapid axon degeneration, our findings with NAM strongly suggest that the cellular dysfunction caused by SARM1 activation can be dissociated from the morphological damage to axons. This distinction is pivotal, as it indicates that SARM1 activation can cause significant neuronal dysfunction without progressing to full-scale degeneration. Recent research has shown that SARM1 activation can occur at sublethal levels across various cell types^8^, including neurons^35–37^. However, the consequences of this sublethal activation on broader cellular function, and whether the toxicity caused by this activation is reversible, have remained largely unexplored. Our study suggests that SARM1 activation significantly affects cellular function even in the absence of axon degeneration in human neurons. Future studies should include experiments assessing the impact of programmed axon death pathway activation on cellular functionality, especially in models where axon degeneration is not observed. This is relevant to numerous preclinical neurodegenerative disease models, especially in human iPSC-derived neuron studies. For instance, most hiPSC-DANs lines derived from PD patients often do not exhibit clear morphological changes. This approach may unexpectedly reveal a broader involvement of programmed axon death in chronic, slow-progressing neurodegenerative disorders, where dysfunction frequently precedes degeneration^57^.

In conclusion, we have comprehensively characterised the detrimental effects of SARM1 activation in human neurons, developed ASOs targeting human SARM1, and demonstrated their effective mitigation of morphological damage and cellular dysfunction. We have demonstrated the reversibility of SARM1 toxicity in human neurons and discussed its significance for clinical applications, both for therapies aiming at inhibiting or activating SARM1. Finally, we highlight the importance of expanding beyond morphology to consider functional impairments, especially in the context of sublethal SARM1 activation. This may reveal key insights for understanding the role of programmed axon death in slow-progressing, chronic neurodegenerative diseases.

## Acknowledgments

We thank members of the Wade-Martins and Coleman laboratories for useful discussions. We also thank Dr Jonathan Gilley for useful comments on the manuscript.

## Author contributions

Conceptualization, AL and KMLC; Methodology, AL, KMLC, and LAM; Validation, AL and KMLC; Formal Analysis, AL and KMLC; Investigation; AL, KMLC, CA, IC, and GO; Resources, AL, LAM, MCC, EDM, DLB, HTZ, MPC, and RW-M; Data Curation, AL and KMLC; Writing - Original Draft; AL and KMLC; Writing - Review & Editing, AL, KMLC, CA, EM, PA-F, HTZ, DLB, GO, MPC and RW-M; Visualization, AL and KMLC; Supervision, AL, MPC, and RW-M; Project Administration, AL; Funding Acquisition, AL, MPC, and RW-M.

## Supplementary figures

**Fig. S1. Validation of the *SARM1^-/-^* line and protection from axotomy-induced axon degeneration.**

**(A)** Representative immunoblots of *SARM1^-/-^* and ISO - CTRL hiPSC-DANs, probed for SARM1, TH and β-actin (loading controls). **(B)** Representative images of axons from *SARM1^-/-^*and ISO - CTRL hiPSC-DANs following axotomy. **(C)** Quantification of the degeneration index for the conditions described in part (B) (mean ± SD; n = 2 from 2 independent differentiations). **(D)** Representative immunofluorescence images of *SARM1^-/-^* and ISO - CTRL hiPSC-DANs stained for TH, MAP2 and DAPI. **(E)** Quantification of differentiation efficiencies of *SARM1^-/-^,* ISO - CTRL hiPSC-DANs and other control lines used in this study (mean ± SD, n = 2, 3 technical repeats from 1 or 2 independent differentiations).

**Fig. S2. Vacor causes dose-dependent neurotoxicity which is blocked by FK866 in hiPSC-DANs.**

**(A)** Representative images of axons from hiPSC-DANs treated with increasing concentrations of vacor or vehicle. **(B)** Quantification of the degeneration index for the conditions described in part (A) (mean ± SEM; n = 4 from 1 independent differentiation; two-way RM ANOVA followed by Tukey’s multiple comparison test; statistical comparison shown is +100 µM Vacor vs +Vehicle). **(C)** Representative images of axons from hiPSC-DANs treated with FK866, vacor or vehicle. **(D)** Quantification of the degeneration index for the conditions described in part (C) (mean ± SEM; n = 4 from 1 independent differentiation; two-way RM ANOVA followed by Tukey’s multiple comparison test; statistical significance shown relative to +100 µM Vacor +100 nM FK866).

**Fig. S3. The strength of protection against axon degeneration correlates with the ASOs’ ability to reduce SARM1 levels.**

**(A)** Representative images of axons from hiPSC-DANs untreated or treated with 15 µM *SARM1* ‘D’ - ASO and Ctrl - ASO following administration of vacor or vehicle. **(B)** Quantification of the degeneration index for the conditions described in part (A) (mean ± SEM; n = 6 from 2 independent differentiations; two-way RM ANOVA followed by Tukey’s multiple comparison test; statistical comparison shown is Ctrl - ASO +100 µM Vacor vs *SARM1* D - ASO +100 µM Vacor). **(C)** Representative images of axons from hiPSC-DANs untreated or treated with 5 µM *SARM1* ‘C’ - ASO and Ctrl - ASO following administration of vacor or vehicle. **(D)** Quantification of the degeneration index for the conditions described in part (C) (mean ± SEM; n = 3 from 1 independent differentiation; two-way RM ANOVA followed by Tukey’s multiple comparison test; statistical comparison shown is Ctrl - ASO +100 µM Vacor vs *SARM1* C - ASO +100 µM Vacor).

**Fig. S4. Mitochondrial impairment begins approximately 2 hours after vacor treatment.**

**(A)** Basal OCR of hiPSC-DANs over time following treatment with vacor or vehicle, normalised to time 0 minutes (mean ± SEM; n = 3 from 1 independent differentiation). **(B)** Basal OCR of *SARM1^-/-^* hiPSC-DANs following treatment with vacor or vehicle, normalised to time 0 minutes (mean ± SD; n = 1, 3 technical repeats from 1 independent differentiation).

**Fig. S5. NAM delays but does not fully block axon degeneration when added hours after vacor treatment.**

**(A)** Schematic overview of the mechanisms by which NAM may reduce vacor toxicity. **(B)** Schematic overview of the experimental design for experiments in (C,D). NAM was added to the media containing concurrently with vacor, or 2 hours and 4 hours after vacor treatment. For this experiment, vacor was still present in the culture media after NAM addition. **(C)** Representative images of axons from hiPSC-DANs following treatments outlined in part (B). **(D)** Quantification of the degeneration index for the conditions described in part (C) (mean ± SEM; n = 6 from 2 independent differentiations; two-way RM ANOVA followed by Tukey’s multiple comparison test; statistical significance shown relative to +100 µM Vacor +1 mM NAM at 4 hr).

