## Supplementary figures and images for "SARM1 activation induces reversible mitochondrial dysfunction and can be prevented in human neurons by antisense oligonucleotides"

### Supplementary figures

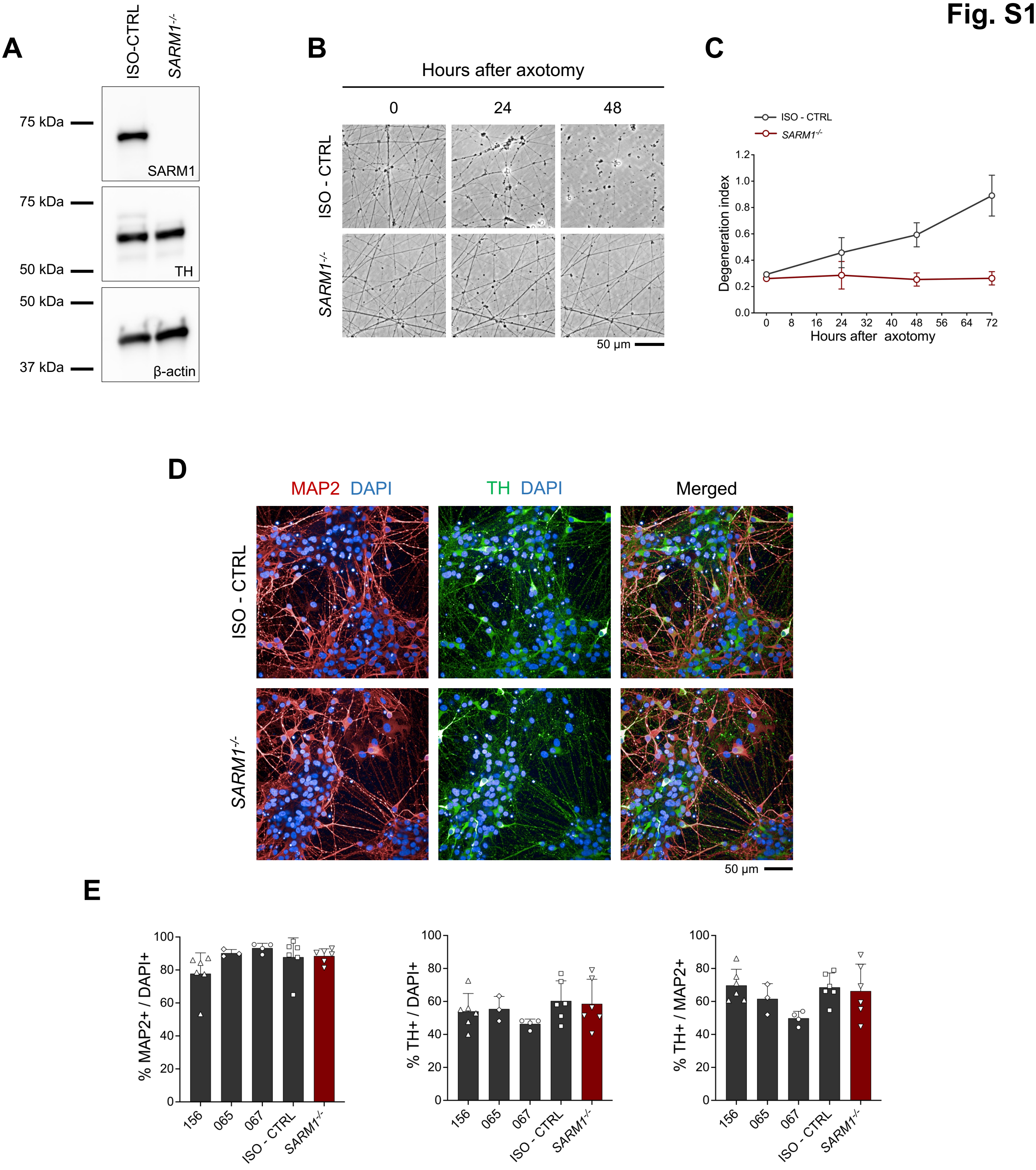

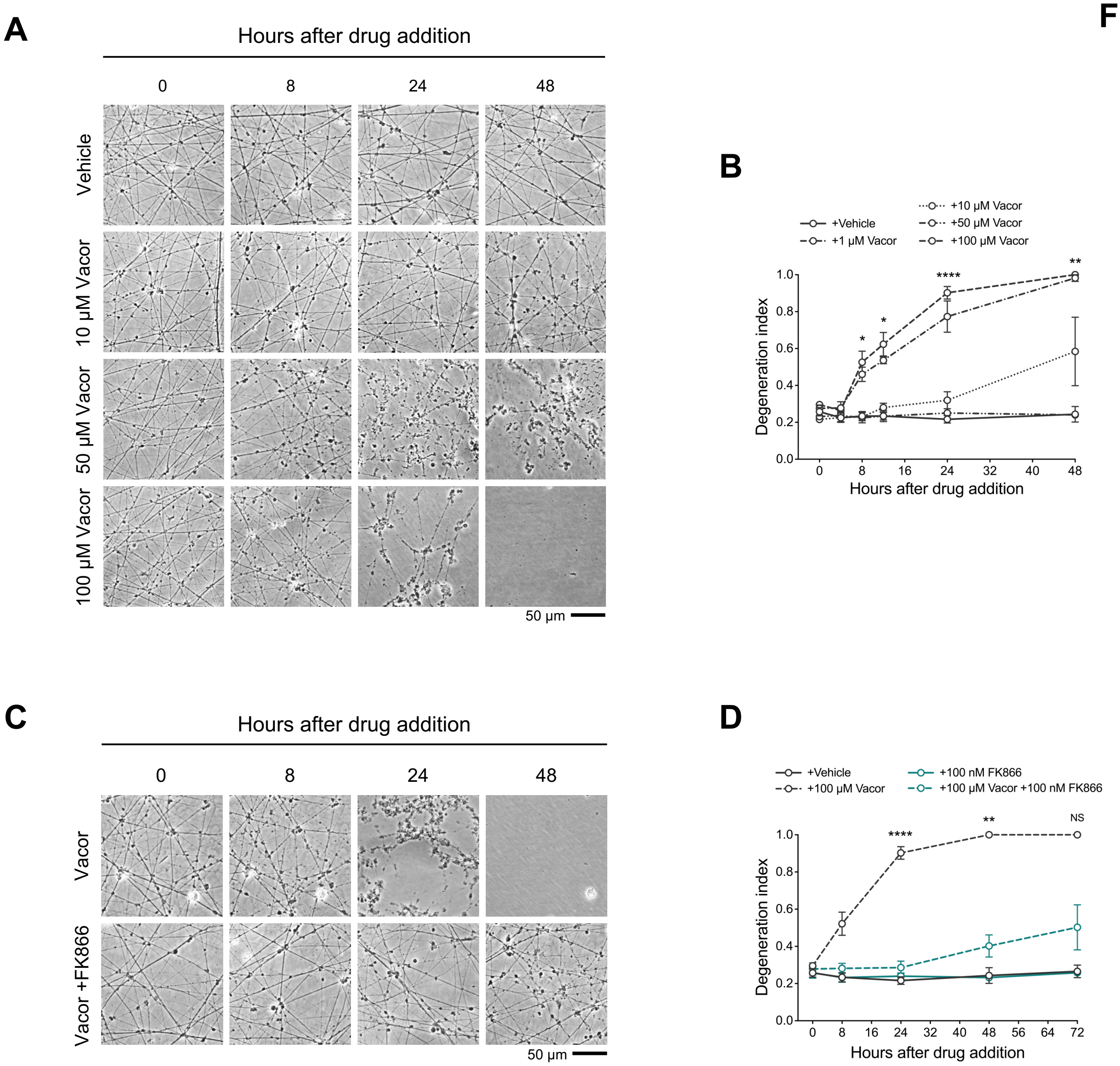

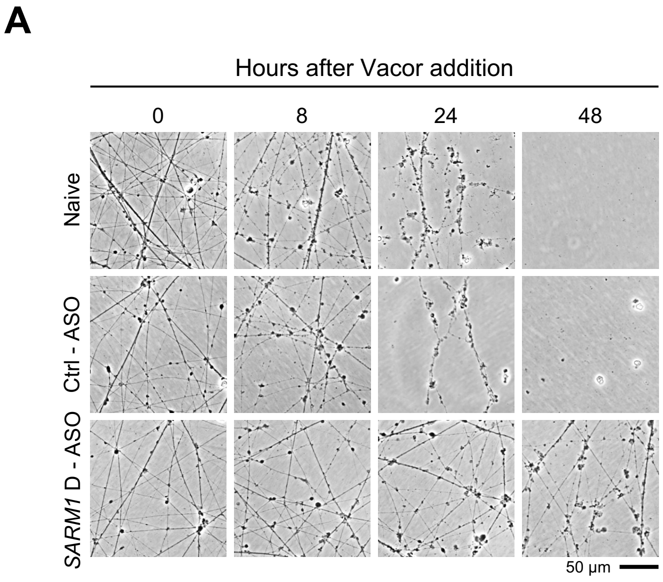

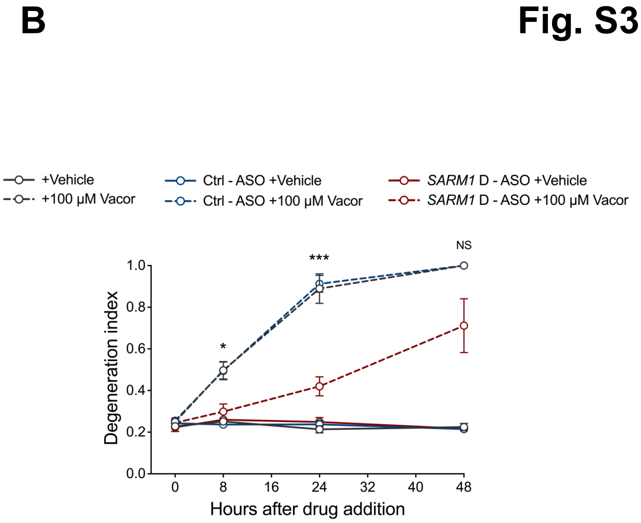

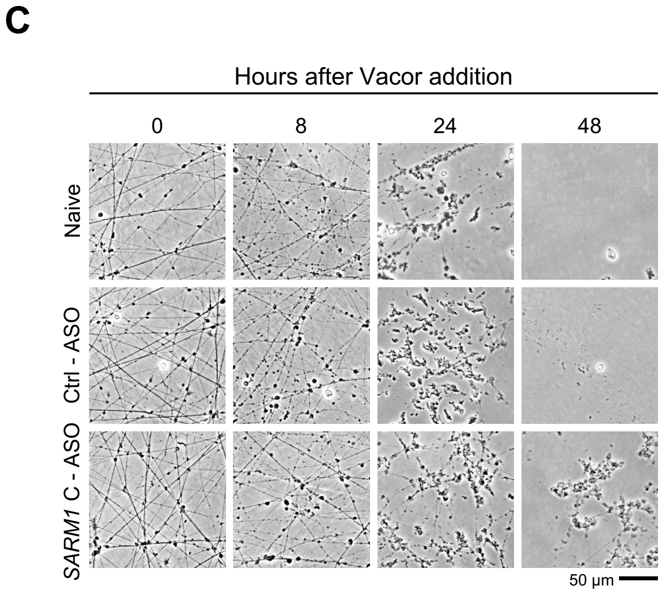

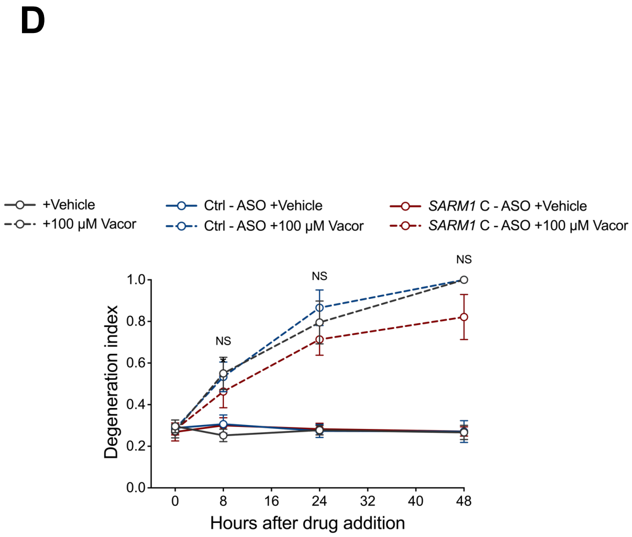

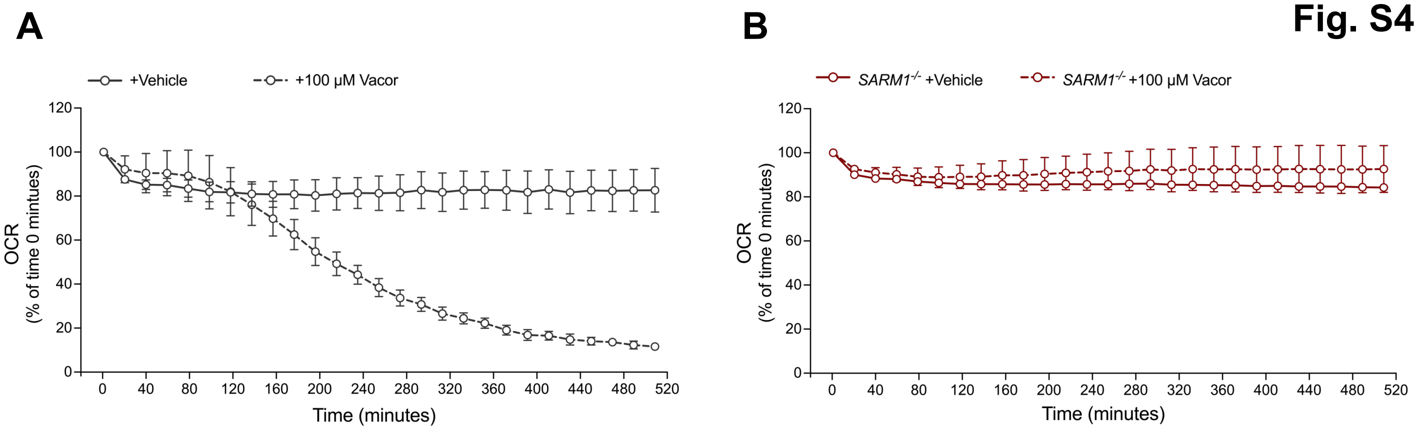

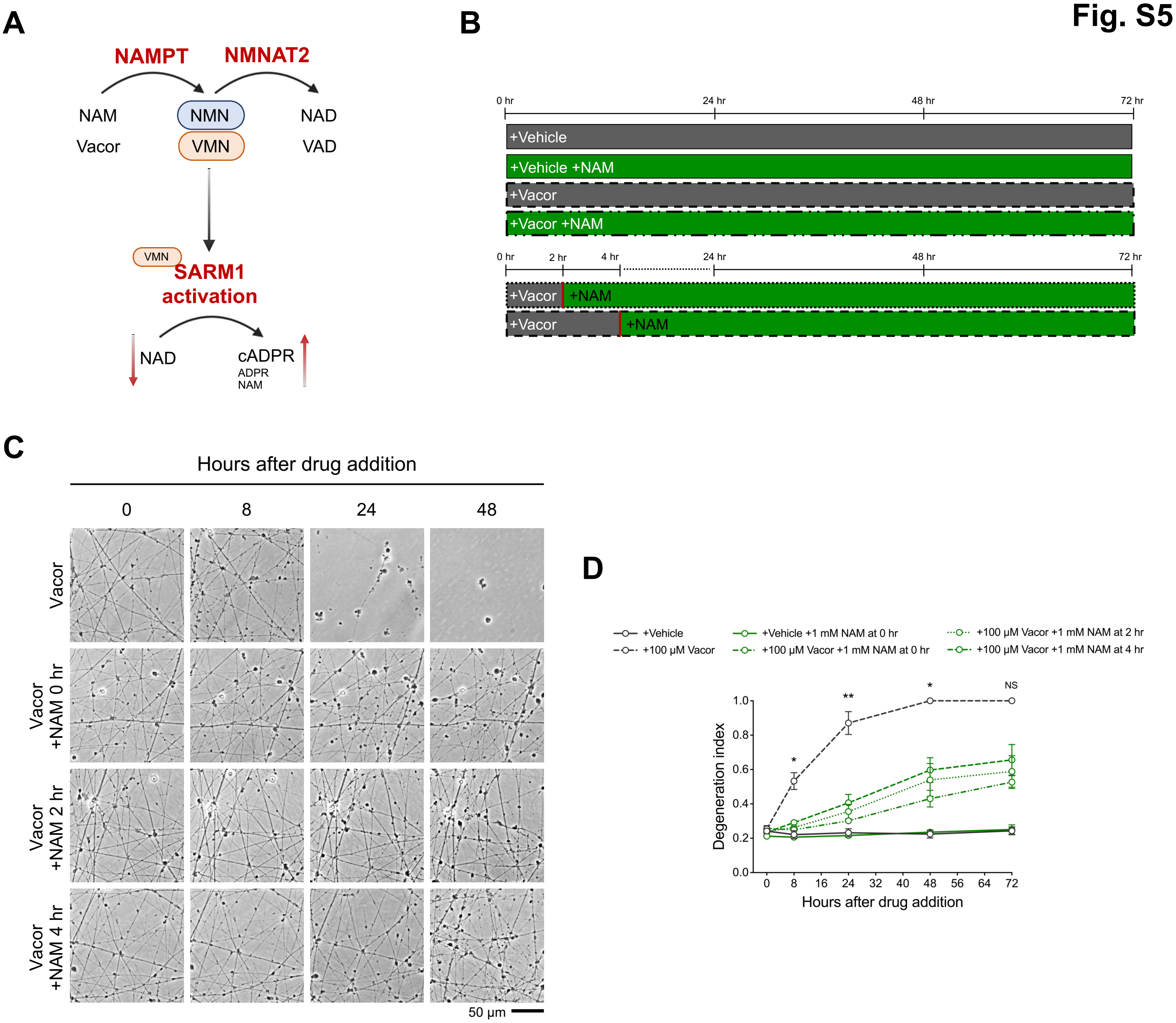
